# Biological characterization of Euscelidius variegatus iflavirus 1

**DOI:** 10.1101/2020.03.31.018010

**Authors:** Sara Ottati, Alberto Persico, Marika Rossi, Domenico Bosco, Marta Vallino, Simona Abbà, Giulia Molinatto, Sabrina Palmano, Raffaella Balestrini, Luciana Galetto, Cristina Marzachì

**Affiliations:** Istituto per la Protezione Sostenibile delle Piante, Consiglio Nazionale delle Ricerche, IPSP-CNR, Strada delle Cacce 73 10135, Torino, Italy; Dipartimento di Scienze Agrarie, Forestali ed Alimentari DISAFA, Università degli Studi di Torino, Largo Paolo Braccini 2, 10095, Grugliasco (TO), Italy

**Keywords:** Leafhopper, phytoplasma vector, insect virus, vertical transmission, horizontal transmission

## Abstract

Virus-based biocontrol technologies represent sustainable alternatives to pesticides and insecticides. Phytoplasmas are prokaryotic plant pathogens causing severe losses to crops worldwide. Novel approaches are needed since insecticides against their insect vectors and rogueing of infected plants are the only available strategies to counteract phytoplasma diseases. A new iflavirus, named EVV-1, has been described in the leafhopper phytoplasma vector *Euscelidius variegatus*, raising the potential to use virus-based application strategies against phytoplasma disease. Here transmission routes of EVV-1 are characterized, and localization within the host reveals the mechanism of insect tolerance to virus infection. Both vertical and horizontal transmission of EVV-1 occur and vertical transmission was more efficient. The virus is systemic and occurs in all life-stages, with the highest loads measured in ovaries and first to third instar nymphs. The basic knowledge gained here on the biology of the virus is crucial for possible future application of iflaviruses as biocontrol agents.

## 1. Introduction

The use of next generation sequencing in the last few years allowed the discovery of many novel viruses in arthropods (Dong et al., 2018; dos Santos et al., 2019; O’Brien et al., 2018). In particular, this technology proved crucial for the identification of covert viruses, those that do not cause overtly pathological effects on their hosts (Nouri et al., 2018). Of the newly discovered virus types, positive-sense RNA viruses are dominant (Bonning, 2020). Among them, iflaviruses (Iflaviridae) form a distinct group in the order *Picornavirales*, (Valles et al., 2017). Effects of some of these viruses include developmental anomalies (Li et al., 2019) behavioural alterations (Wells et al., 2016), tissue pathologies (Brettell et al., 2017), and premature death (Geng et al., 2017) of the insect host, while others do not cause any evident symptoms (dos Santos et al., 2019; Murakami et al., 2013). Transmission modalities of these viruses have been explored, leading to a multifaceted infection model. Horizontal transmission (oral route) is the most efficient means of spread of some species, for example Nilaparvata lugens honeydew virus-1 of the brown leafhopper (Murakami et al., 2013), Helicoverpa armigera iflavirus (Yuan et al., 2017), and Nora Virus of the cotton bollworm (Yang et al., 2019), and Pyrrhocoris apterus virus 1 in red firebugs (Vinokurov and Koloniuk, 2019). In contrast, Antheraea pernyi Vomit Disease of Chinese oak silkmoth (Geng et al., 2017) and Spodoptera exigua iflaviruses-1 and 2 of the beet armyworm (Virto et al., 2014) appear to be primarily spread by vertical transmission. The Deformed Wing Virus (DWV) of the honeybee is transmitted both horizontally and vertically, and by *Varroa destructor* mite as vector (Chen et al., 2006; Yue and Genersch, 2005).

The genomic features, phylogenetic analysis, and prevalence of a new member of the family *Iflaviridae*, Euscelidius variegatus virus 1 (EVV-1), were previously reported (Abbà et al., 2017) as an additional result obtained during the *de novo* assembly of transcriptome from the hemipteran leafhopper *Euscelidius variegatus* Kirschbaum (Galetto et al., 2018). EVV-1 did not induce any evident symptoms in the *E. variegatus* population surveyed but, when laboratory-reared, produced infections of 100% prevalence (Abbà et al., 2017). EVV-1 forms non-enveloped, icosahedral particles, and the ssRNA(+) genome encodes a single polyprotein of 3132 amino acids, which is post-translationally processed.

*Euscelidius variegatus* (Cicadellidae: Deltocephalinae) is a palearctic, multivoltine and polyphagous species, widespread in Europe and North America. This species is economically important because it is a natural vector of ‘*Candidatus* Phytoplasma asteris’ (chrysanthemum yellows strain) and a laboratory vector of the Flavescence dorée phytoplasma (reviewed in EFSA Panel on Plant Health (PLH), 2014. Phytoplasmas are plant-pathogenic bacteria that cause severe symptoms in affected plants, leading to heavy economic losses of many crops worldwide (Tomkins et al., 2018). In particular, Flavescence dorée phytoplasma of grapevine is a quarantine pathogen, which strongly limits European viticulture (EFSA Panel on Plant Health (PLH), 2014). Phytoplasmas are obligate parasites with a dual life cycle, infecting phloem of host plant as well as the body of the insect vector. Several lines of evidence indicate that interactions between phytoplasmas and vector host cells are very strict and may regulate transmission ability (reviewed in Rossi et al., 2019). Molecular relationships between pathogens and insect hosts contribute to determining phytoplasma transmission specificity and plant susceptibility, as well as feeding behaviour, ecological dispersal and plant host range vectors (Bosco and D’Amelio, 2010). Insects play a key role in phytoplasma epidemiology and, therefore, the main control strategies to limit these pathogens are insecticide treatments against the vector species (Tomkins et al., 2018).

The urgent need for more targeted and sustainable pest management approaches in fighting phytoplasma diseases includes the exploration for alternative biocontrol agents such as insect viruses. Many fundamental biological features of EVV-1 were unknown, impairing potential applications of this agent in biocontrol strategies or as a virus-induced gene silencing (VIGS) vector. To fill the gap of knowledge about EVV-1 biology, we investigated the presence of EVV-1 virus in different developmental stages and organs of *E. variegatus*, as well as both horizontal and vertical transmission of this iflavirus among insect populations.

## 2. Materials and methods

### 2.1 Insects

*Euscelidius variegatus* in the Turin laboratory colonies were originally collected in Piedmont Italy and reared on oats, *Avena sativa* (L.), inside plastic and nylon cages in growth chambers at 20–25 °C, L16:D8 photoperiod. A French population of *E. variegatus* was kindly provided by Dr. Xavier Foissac (INRA, UMR1332 Biologie du Fruit et Pathologie, Bordeaux, France) and reared in Turin laboratory under the above described conditions. The original Turin colony (EvaTo), was naturally infected with EVV-1 virus with a 100% prevalence, whereas the French colony (EvaBx) appeared to be EVV-1 free, according to molecular detection and electron microscopy observation (Abbà et al., 2017). The two colonies were routinely confirmed to be EVV-1 infected (EvaTo) and free (EvaBx) by RT-qPCR, as described below.

### 2.2 Collection of different insect life stages and organ dissection

To determine the viral loads in different insect life stages, total RNAs were extracted from laid eggs, I-V instar nymphs and adults of virus infected EvaTo population. Preliminary experiments showed that egg collection was easier when eggs were laid on *Arabidopsis thaliana* plants than on *A. sativa* or *Vicia faba*. Therefore, approximately 100 EvaTo female adults were caged on 15 *A. thaliana* plants for 4 d and then eggs were collected with sterile forceps using a stereomicroscope. Seven pooled samples, each containing approximately 30 eggs, were obtained. To exclude surface viral contamination, egg samples were sterilized according to Prado et al., (2006) and then stored at −80°C prior to RNA extraction. For each nymphal stage, 10 pooled samples (each with three EvaTo specimens) were collected and stored at −80°C until RNA extraction. Newly emerged EvaTo adults (five males and five females) were also collected and stored.

To estimate the viral presence in different insect organs, total RNAs were extracted from Malpighian tubules, salivary glands, guts, ovaries, testes and hemolymph of newly emerged EvaTo adults. Three hemolymph samples (each with hemolymph collected from 10 specimens) were obtained by removing the head of a CO_2_-anaesthetized adult and drawing 0.5 μL hemolymph from the inner thorax cavity with a Cell Tram Oil microinjector (Eppendorf), using a stereomicroscope. For the remaining organs, three pooled samples (each made of five dissected organs) were obtained. Organs were carefully separated with forceps and needles using a stereomicroscope, rinsed with phosphate-buffered saline (PBS) solution and stored at −80°C until RNA extraction.

### 2.3 Virus vertical transmission route test

To determine whether EVV-1 is vertically transmitted, virus free EvaBx adults were injected with a fresh extract of EvaTo individuals. Microinjection was chosen as infection modality to maximize the probability of obtaining a high number of infected EvaBx adults. These were then used as F0 parents to test vertical transmission to the offspring (F1 and F2 generations). The fresh extract containing virus particles was prepared by crushing 30 EvaTo adults in 900 μL of cold filter-sterilized injection buffer (300 mM glycine, 30 mM MgCl2, pH 8.0). The extract was clarified by slow centrifugation (10 min, 800 g), and passed through a 0.22 μm pore-size filter. All extraction steps were done at 4°C. Newly emerged EVV-1 free EvaBx adults were CO_2_ anaesthetized and injected with 0.5 μL of the virus-containing extract between two abdominal segments using a stereomicroscope with a Cell Tram Oil microinjector (Eppendorf). As negative control in order to exclude any procedural contamination, a group of EvaBx insects was injected with injection buffer only. Injected insects were isolated on fresh oat plants and sampled at 4 and 10 days post injection (dpi). The F1 and F2 adults were collected at 60 and 120 dpi, respectively. All collected insects (F0, F1 and F2) were stored at −80°C until RNA extraction to check the presence of EVV-1. The experiment was repeated twice.

### 2.4 Virus horizontal transmission route test

To determine whether EVV-1 is horizontally transmitted, EVV-1 free EvaBx adults were caged together with infected EvaTo. In order to distinguish individuals belonging to the two original populations, three strategies were carried out: EvaBx third instar nymphs caged with EvaTo adults (Experiment 1); EvaBx female adults caged with EvaTo male adults (Experiment 2); and EvaBx male adults caged with EvaTo female adults (Experiment 3). For each experiment, 50 EvaBx insects were co-fed on the same oat plant together with 50 EvaTo individuals for two weeks, then transferred to a new plant in a new clean and sampled three weeks later for virus detection. Each experiment was repeated twice.

After the positive results of co-feeding, three possible horizontal transmission modalities were specifically tested i) fecal-oral route, ii) virus infection via plant, and iii) cuticle penetration. Fecal-oral transmission modality was assessed according to Murakami et al. (2013). About 60 EvaTo adults were confined in a 50-mL conical tube for 1 h. The insects were then removed from the tube, and the excreted honeydew was collected by rinsing the wall of the plastic tube with 4 mL of artificial feeding medium (5% sucrose, 10 mM Tris/Cl, 1 mM EDTA, pH 8.0. An aliquot (400 μL) of this solution was stored at −80°C for RNA extraction to confirm the presence of the virus, while the remaining solution was used to artificially feed EvaBx adults. To this purpose, non-viruliferous insects were confined within small chambers. Six small cages were set up, each with five EvaBx insects allowed to feed for 48 h on 600 μL of honeydew solution laid between two layers of stretched Parafilm. As negative controls, three small cages were set up with artificial medium only, devoid of honeydew. Following artificial feeding, live insects were caged on oat plants for 10 d (Murakami et al., 2013), then collected and stored at −80°C, until RNA extraction for virus detection.

To assess the plant-mediated horizontal transmission of EVV-1, virus presence was tested in *A. sativa* plants exposed to either EvaTo or EvaBx insects, following leaf surface sterilization or no sterilization (Prado et al., 2006). Virus transmission through the host plant was tested by isolating about 50 EvaTo adults for 1 week on an oat plant. The virus-positive insects were then removed, while the oat plant was used to feed about 50 EvaBx adults for the following week. EvaBx adults were then transferred onto a fresh oat plant and five were tested for the presence of EVV-1 every 7 d for 1 mo.

To assess EVV-1 ability to penetrate through the insect cuticle, EvaBx nymphs were submerged into fresh virus extract. For this experiment, second instar nymphs were used, as the highest virus level was detected at this life stage in EvaTo and the insect dimensions allowed an easier manipulation compared to the smaller and more delicate first instar nymphs. Briefly, EvaBx nymphs were CO_2_ anaesthetized and submerged for 5 min in 300 μL fresh virus solution prepared by crushing ten EvaTo adults in 400 μL of PBS with a cocktail of protease inhibitors added (Pierce Protease Inhibitor Mini Tablets, EDTA-Free, Thermo Scientific). The extract was clarified by slow centrifugation (5 min, 4500 g). Submerged nymphs were then maintained on an oat plant and collected as adults for RNA extraction and virus detection.

### 2.5 RNA extraction, cDNA synthesis, EVV-1 qPCR assays and statistical analyses

Total RNAs were extracted from samples of single *E. variegatus* adults, and pooled samples of i) 30 eggs, ii) three nymphs, iii) three dissected organs, iv) hemolymph collected from 10 insects, v) honeydew collected from 60 insects and vi) 500 mg of oat leaves, with Direct-zol RNA Mini Prep kit (Zymo Research), following manufacturer’s protocol. For each sample, cDNA was synthesized from total RNA (1 μg) with random hexamers using a High Capacity cDNA reverse transcription kit (Applied Biosystems).

A multiplex qPCR assay was developed to detect and quantify virus presence in *E. variegatus* adults. To detect EVV-1, primers and FAM-labelled TaqMan probe were designed on the virus capsid 1 coding sequence (Evv1Cap1Fw 5’-GACCATTATCGCGCTAATG-3’, Evv1Cap1Rv 5’-AGTGCTCATCATAGGACA-3’, Evv1Cap1Probe 5’-FAM-ATTCTCGTAGCCAACTGCCAAAC-3’). To check that all the cDNA samples were properly amplifiable, the *E. variegatus* glyceraldehyde-3-phosphate dehydrogenase (GAPDH) transcript, was chosen as endogenous control (GapFw632 5’-ATCCGTCGTCGACCTTACTG-3’, GapRv819 5’-GTAGCCCAGGATGCCCTTC-3’, GapEvProbe 5’-HEX-ATATCAAGGCCAAGGTCAAGGAGGC-3’). Standard conditions were used for qPCR assay and detailed in Supplementary Method File. Mean virus copy numbers were used to express virus amount as EVV-1 copy numbers/insect GAPDH transcript.

For comparison of viral load in different insect life stages and organs, virus load was expressed as EVV-1 copy numbers/ng of cDNA, as GAPDH transcript levels varied among the different organs and life stages (Supplementary Tables S1 and S2).

SigmaPlot version 13 (Systat Software, Inc., www.systatsoftware.com.) was used for statistical analyses. To compare viral load among different categories (life stages, organs and vertical transmission experiments) ANOVA, followed by the proper post hoc test, or Kruskal Wallis test, when the parametric analysis assumptions were not met, were used. To compare viral load between two groups of samples (male vs. female EvaTo adults, virus exposed EvaBx vs. EvaTo samples in horizontal transmission experiments) t-test was used.

## 3. Results

### 3.1 EVV-1 qPCR diagnostic and quantitative assay

A multiplex qPCR assay was optimized to detect and quantify the EVV-1 virus in *E. variegatus* adults, using primers and TaqMan probes targeting viral EVV-1 capsid 1 and insect GAPDH coding sequences. The viral genomic portion coding the virus capsid 1 protein was selected to maximize the specificity of EVV-1 detection, as this genomic portion was poorly conserved among other iflaviruses according to BLAST analysis (not shown). Standard curves of plasmids harbouring viral and insect target amplicons showed 96.5 and 103.1% qPCR efficiencies, respectively, with a 0.998 R^2^ for both curves. The most diluted standard point (10^2^ copy numbers/μL) was detected before the 35^th^ amplification cycle by both viral and insect primer sets. The test successfully detected EVV-1 in single EvaTo individuals (n=10), whereas no amplification was obtained from any of the 10 tested EvaBx. Quantification of EVV-1 in the EvaTo adults ranged from 0.21 to 0.84 EVV-1 copy numbers/insect GAPDH transcript.

### 3.2. EVV-1 distribution in different insect life stages and organs

EVV-1 was detected in the eggs and all insect developmental stages (I-V instar nymphs and adults) of the naturally infected EvaTo population. Male and female EvaTo individuals were considered together for the analyses, as no significant differences in viral loads were found (t-test t=1.448, P=0.186) (Supplementary Table S1). Significant differences were found among developmental stages (ANOVA F=10.367, P<0.001) and I, II, III instar nymphs showed the highest viral loads, as indicated by pairwise multiple comparison according to Duncan’s multiple range test (Fig. 1 and Supplementary Table S1).

**Figure 1.**
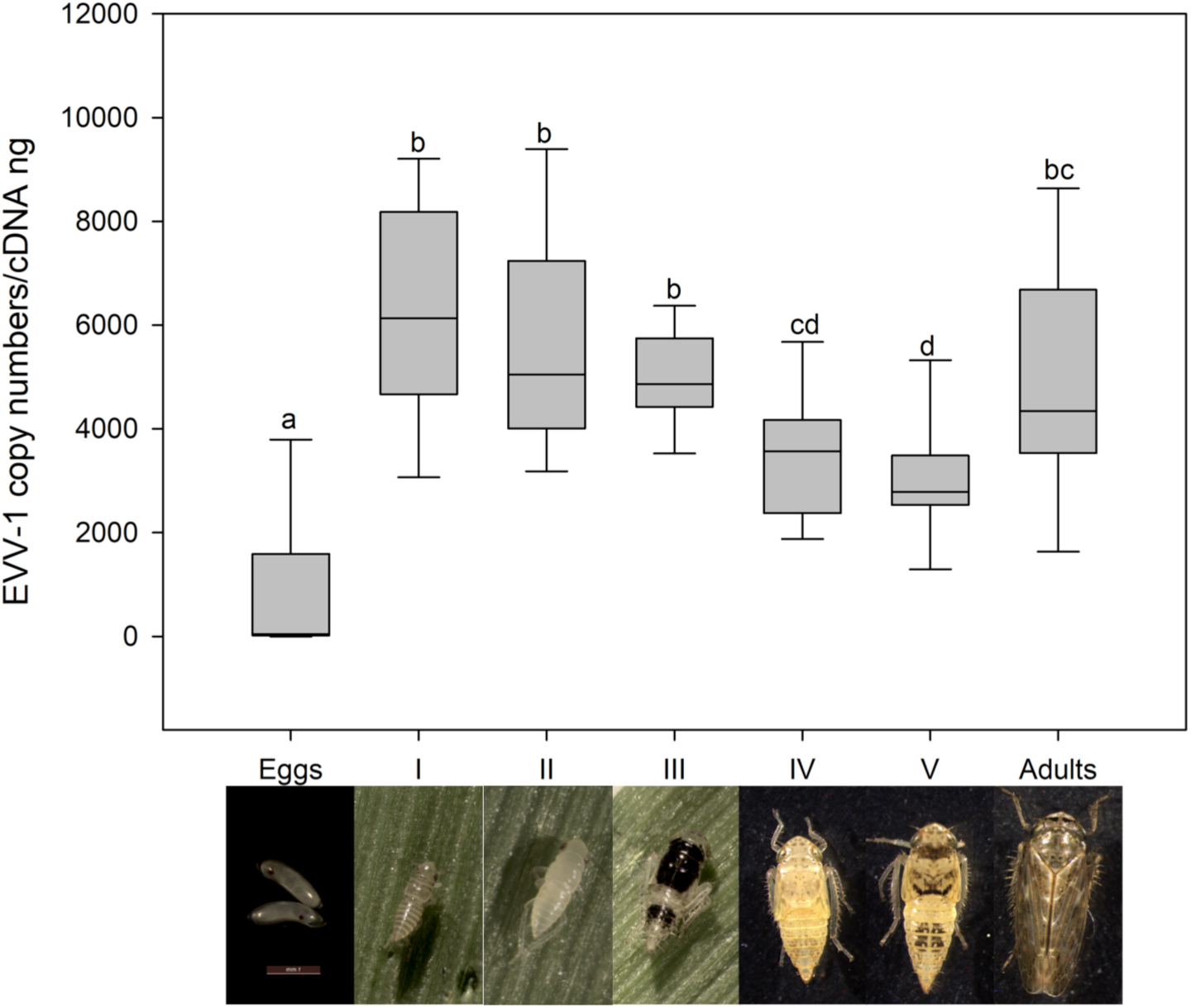
Euscelidius variegatus virus 1 (EVV-1) load in different host life stages. Different letters indicate significant differences in mean viral load measured in the sample groups. First to fifth instar nymphs are indicated by Roman numerals and different life stages are depicted beneath the graph.

EVV-1 was detected in all collected insect organs (Malpighian tubules, salivary glands, guts, ovaries, testes, and hemolymph) of the EvaTo population. Viral loads ranged from 8606 in the ovaries to 1 EVV-1 copy numbers/ng of cDNA in the testes. The high viral load found in ovaries significantly differed from that found in the testes (ANOVA on Ranks, Kruskal-Wallis test H=15.415, P=0.009), (Fig. 2 and Supplementary Table S2).

**Figure 2.**
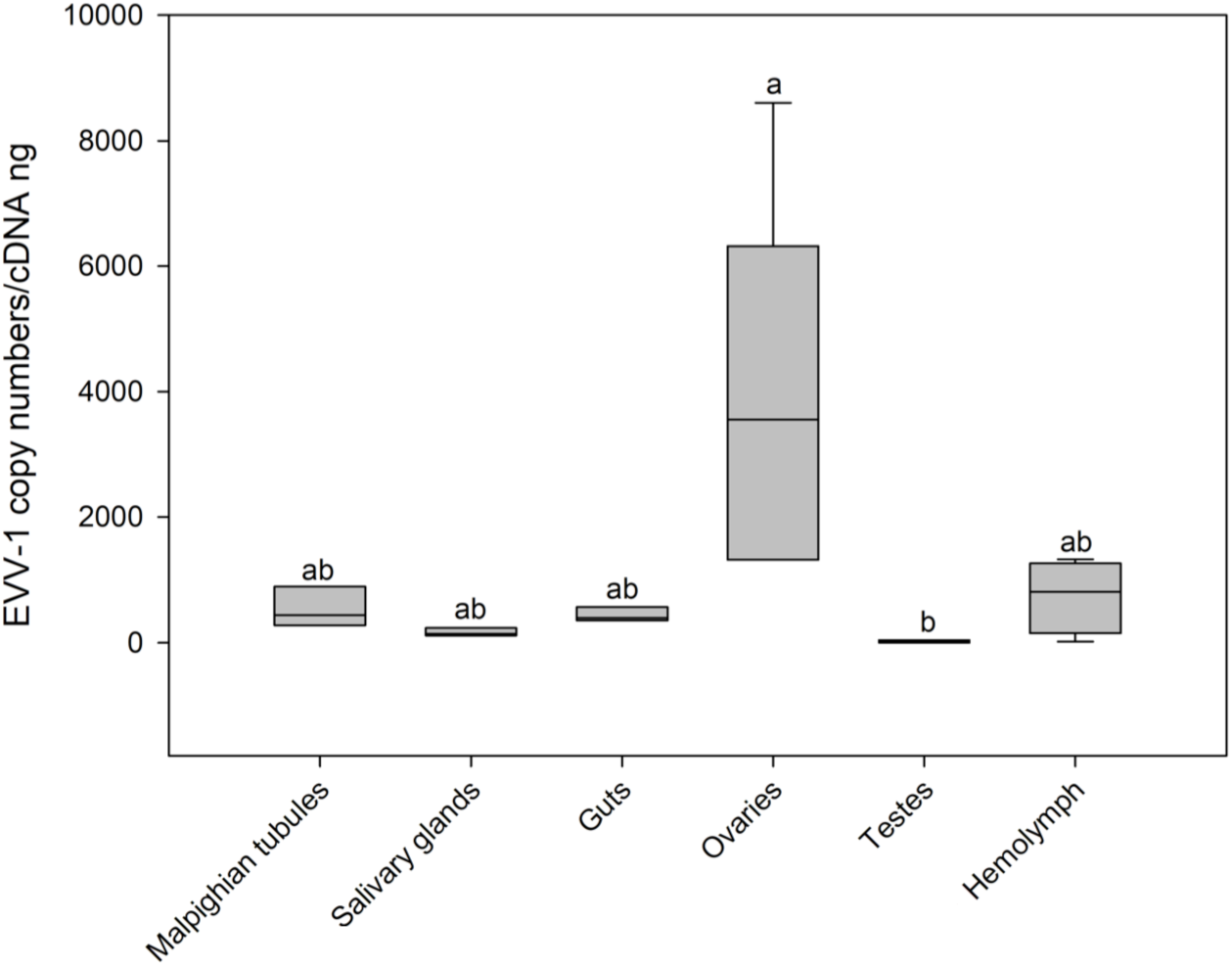
Euscelidius variegatus virus 1 (EVV-1) load in different insect organs. Different letters indicate significant differences in mean viral load measured in the sample groups.

### 3.3 Vertical transmission

Virus injection was selected as the most effective infection modality to obtain a high number of newly infected EvaBx individuals (F0) and test their ability to transmit the virus to progeny (F1 and F2). Virus presence was detected in 95 and 100 % injected insects sampled at 4 and 10 dpi, respectively, as well as in all the F1 and F2 tested insects (Supplementary Table S3), demonstrating that vertical viral transmission occurs with very high frequency. Interestingly, mean EVV-1 loads measured in F1 and F2 samples were significantly higher than those detected in injected F0 insects sampled at 4 dpi and in EvaTo adults (ANOVA on Ranks, Kruskal-Wallis test H=22.769, P<0.001) (Fig. 3). The virus was detected in EvaTo fresh extract injected in EvaBx specimens, while no amplification was obtained from EvaBx adults injected with buffer only (negative control).

**Figure 3.**
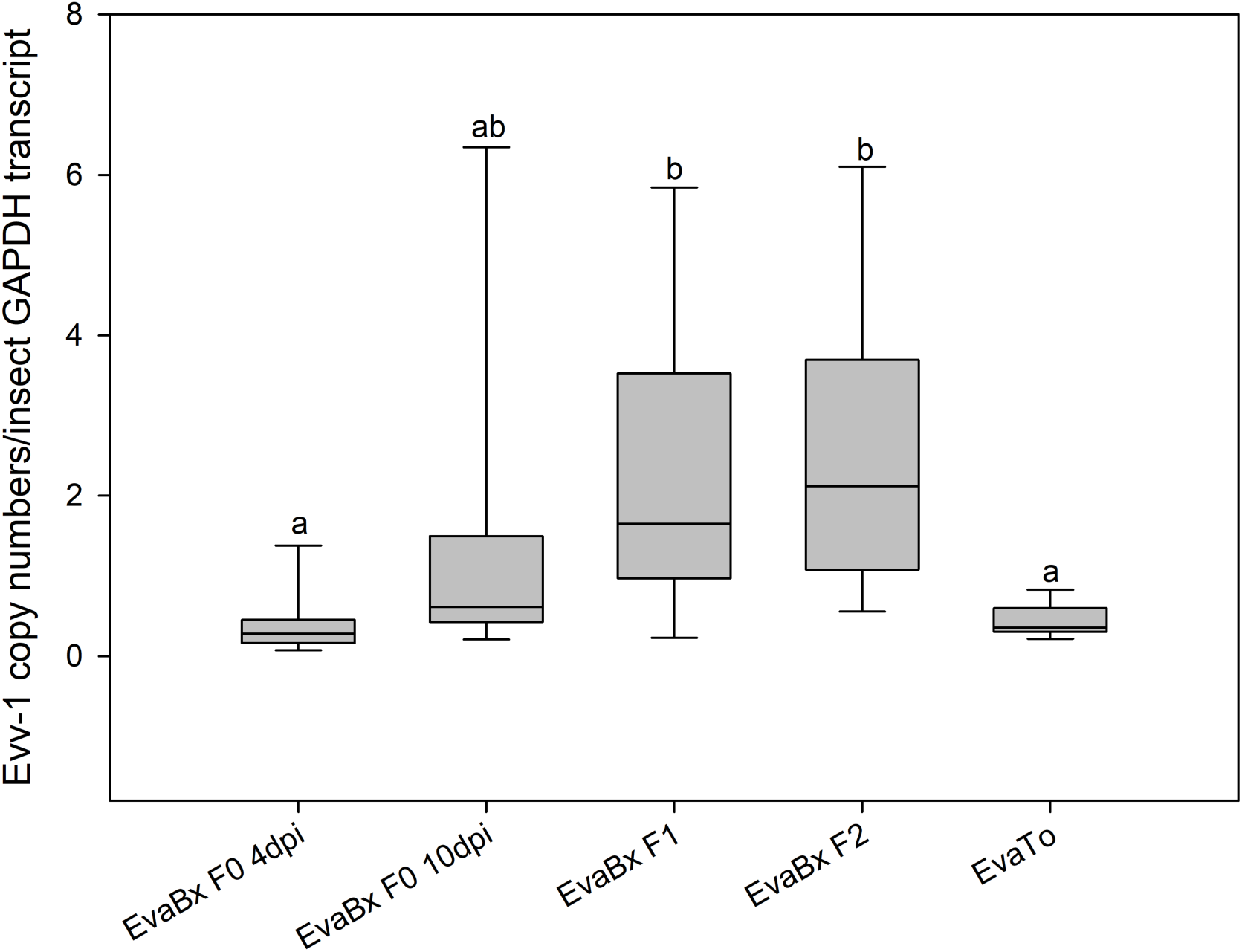
Vertical transmission of Euscelidius variegatus virus 1 (EVV-1) injected into uninfected *Euscelidius variegatus* (EvaBx). Different letters indicate significant differences in mean viral load measured in the sample groups. Days post injection (dpi) indicate the sampling date after viral injection. Viral load measured in infected colony *Euscelidius variegatus* (EvaTo) adults was used for comparison.

### 3.4 Horizontal transmission

The co-feeding experiment, in which EvaTo insects were isolated together with EvaBx specimens on the same plant, indicated that horizontal viral transmission may occur. The overall mean rate of virus transmission was 8.0% (6/75), comprehensively considering the three different co-feeding experimental approaches (indicated in Table 1). The mean viral load, measured in the infected EvaBx insects three weeks after co-feeding with EvaTo, was 0.23 EVV-1 copy numbers/insect GAPDH transcript ± 0.10 (SEM, N=6), with values ranging from 6.27E-04 to 5.94E-01. The mean viral load measured in these virus-exposed EvaBx samples did not significantly differ from that measured in EvaTo adults, which was 0.44 EVV-1 copy numbers/insect GAPDH transcript ± 0.14 (SEM, N=10) (t-test t=1.815, P=0.091).

**Table 1.** Detection and quantification results of horizontal transmission of Euscelidius variegatus virus 1 (EVV-1) to uninfected colony EvaBx. Mean quantification cycles (Cq) obtained from insect glyceraldehyde-3-phosphate dehydrogenase (GAPDH) qPCR assay confirmed that all samples were amplifiable.

| **Virus** |  | **N° EVV-1** | **Mean EVV-1** | **Mean Cq ± SEM (N)** |
| --- | --- | --- | --- | --- |
| **infection** | **Sample description** | **positive/analysed** | **viral load\* ±** | **of GAPDH assay** |
| **modality** |  | **samples** | **SEM (N)** |  |
| Co-feeding | EvaBx nymphs +<br>EvaTo adults | 3/47 | 0.27 ± 0.11 (3) | 18.93 ± 0.36 (47) |
|  | EvaBx females +<br>EvaTo males | 2/17 | 2.56E-03 ±<br>1.94E-03 (2) | 23.45 ± 0.36 (17) |
|  | EvaBx males +<br>EvaTo females | 1/11 | 0.59 (1) | 19.44 ± 0.84 (11) |
| Fecal-oral<br>route | EvaBx adults fed on<br>EvaTo honeydews | 0/12 | / | 20.30 ± 0.28 (12) |
| Plant-mediated<br>route | EvaBx adults fed on plant<br>exposed to EvaTo adults | 0/20 | / | 20.79 ± 0.44 (20) |
| Cuticle<br>penetration | EvaBx nymphs submerged<br>in EvaTo extract | 10/105 | 0.68 ± 0.35 (10) | 17.85 ± 0.16 (105) |

\*Viral load expressed as EVV-1 copy numbers/insect GAPDH transcript.

When exploring the possible routes of EVV-1 horizontal transmission, none of the EvaBx insects artificially fed on virus-infected honeydew solution were positive for EVV-1 presence (Table 1). Although the presence of EVV-1 in EvaTo honeydew was confirmed by qPCR, the possibility of virus transmission via fecal-oral route was not confirmed under our experimental conditions.

None of the 20 analysed EvaBx adults tested positive for EVV-1 presence (Table 1) following isolation on *A. sativa* plants that were exposed to EvaTo insects, thus excluding the plant as efficient transmission route. Indeed, while EVV-1 was detected only in one of six samples of *A. sativa* exposed to EvaTo insects, the virus was absent from all the six analysed plants following surface sterilization of the leaf before RNA extraction, indicating that viral contamination of the leaves may occur. The absence of EVV-1 in plants exposed to EvaBx insects was also confirmed, as expected (0 positive samples out of six total analysed samples).

Viral transmission through insect cuticle of second instar EvaBx nymphs occurred in 9.5% of the treated insects (Table 1). The viral loads measured in these infected EvaBx insects ranged from 0.09 to 1.79 EVV-1 copy numbers/insect GAPDH transcript. No significant differences were found between the mean viral loads measured in these virus-exposed EvaBx samples and those measured in EvaTo adults (t-test t=-1.414, P=0.174).

## 4. Discussion

We studied the distribution and transmission routes of the Euscelidius variegatus iflavirus 1 and showed that infection with the virus is systemic and occurs in all life stages. As well, it is vertically transmitted to offspring with high efficiency. Elucidating these biological parameters is a prerequisite to exploring the potential application of iflaviruses as biocontrol agents of phytoplasma vectors. Our work also provides information needed to exploit iflaviruses as molecular tools to study insect genes involved in pathogen transmission through the construction of infectious viral clones.

Iflaviruses colonize their arthropod hosts in different ways. In some cases, viruses do not exhibit tropism (restriction to specific organs or tissues). Deformed Wing Virus (DWV) can be detected in all parts of the bee body (Martin and Brettell, 2019), and Antheraea pernyi Iflavirus (ApIV) in the head, epidermis, hemocytes, gut, fat body, ovary and testis of the Chinese oak silkmoth, ApIV (Geng et al., 2017). On the contrary, other iflaviruses show some organ tropism, such as the Nora Virus of the cotton bollworm which show higher loads in midgut tissues (Yang et al., 2019) or the Helicoverpa armigera iflavirus, which shows higher levels in fat bodies compared to other insect body tissues. Molecular analysis showed that EVV-1 is present in different *E. variegatus* tissues and organs as well as in different life stages (from egg to adult). The highest viral load was detected in I-III instar nymphs and may indicate that these stages support the peak of viral replication, even though northern blot analysis did not detect an accumulation of negative strands in nymphal stages nor in adults (Abbà et al., 2017). The presence of detectable replicative intermediates may be limited in time and difficult to detect. In line with our results, increase of viral load over time was observed in pupae of the Chinese oak silkmoth following the injection of ApIV purified particles (Geng et al., 2017). The high EVV-1 viral load measured in the ovaries suggests a key role of this organ in the viral replication cycle. Indeed, this feature may be common to iflaviruses infecting Hemiptera. For example, like EVV-1, the Laodelphax striatellus Iflavirus 1 (LsIV1) is present in eggs and ovaries of the small brown planthopper (Wu et al., 2019). Infection data from different iflaviruses seems, therefore, to delineate an assorted model of insect body colonization, that requires case-by-case characterization.

Under our experimental conditions, EVV-1 was efficiently transmitted to the offspring. Indeed, following injection of virus-free parents, 100% of the F1 and F2 offspring were infected. The development of an efficient multiplex fluorescent qPCR assay, based on glyceraldehyde-3-phosphate dehydrogenase (GAPDH) as an endogenous gene for normalization of quantitative data and as internal control of cDNA quality, allowed the quantitative detection of EVV-1, a prerequisite to characterize its distribution and transmission modalities. Interestingly, the mean viral load of the F1 and F2 individuals was significantly higher than that of naturally infected EvaTo adults. This result supports the hypothesis of a higher viral replication rate in the experimentally infected insects than in the naturally infected insects. The absence of evident fitness costs was in line with preliminary results showing that longevity and fecundity were not negatively affected by EVV-1 infection. Similarly, in several mosquito species, a state of tolerance response against covert arboviral infections has been described leading to persistent infection in the host with little fitness cost (Goic et al., 2016).

Horizontal transmission also has been reported for some iflaviruses (Murakami et al., 2013; Yuan et al., 2017), and occasionally occurred for EVV-1 under our experimental conditions, although less efficiently than vertical route. While Nilaparvata lugens iflavirus 1 (NHLV-1) is efficiently transmitted to virus-free brown leafhoppers by feeding on infected honeydew (Murakami et al., 2013), the EVV-1 virus was not transmitted to EvaBx individuals by feeding, despite EVV-1 presence in EvaTo honeydew added to the artificial diet. This suggests that EVV-1 is not able to cross the gut barrier. EVV-1 could be sporadically detected in unsterilized leaves from plants exposed to virus-infected insects, probably due to honeydew contamination, but it was never detected after sterilization of leaf surface. The phloem feeding habit of *E. variegatus* may explain its inability to acquire EVV-1 through the host plant.

Silencing of insect genes by RNA interference is a promising approach to control insect pests, and it has been applied to several Hemipteran species including aphids (Yu et al., 2016), psyllids (Lu et al., 2019), and whiteflies (Shi et al., 2019). Efficient delivery of RNA interfering molecules is difficult to achieve for phloem feeders, and microinjection (Mutti et al., 2006) or plant-mediated silencing strategies (Jaubert-Possamai et al., 2007; Pitino et al., 2011) have been explored. In particular, soaking of nymphs of the psyllid *Diaphorina citri*, vector of ‘*Candidatus* Liberibacter solanacearum’, in a solution containing dsRNA silencing molecules provides efficient silencing of different target genes (Killiny et al., 2014; Yu et al., 2017). EVV-1 was transmitted to nymphs following their immersion in a solution containing the virus, suggesting that the viral particles are able to penetrate the nymphal cuticle or to pass through the tracheae and then replicate in the insect body. This soaking-based strategy of viral delivery may be further explored as a high-throughput technique to increase the horizontal transmission efficiency of phloem feeder viruses. Also, the ability of EVV-1 to penetrate the nymphal cuticle or the tracheae may also explain the occasional transmission of the virus to some EvaBx individuals, following co-feeding together with virus infected EvaTo individuals. In conclusion, both vertical and cuticle/tracheal horizontal transmission routes may explain the 100% EVV-1 prevalence observed in the mass reared EvaTo lab colony.

Although the idea behind the application of entomopathogens as biocontrol agent dates back to the nineteenth century (Metchnikoff, 1880), the use of insect-specific viruses in plant protection is lately gaining attention. Iflaviruses may infect their hosts and transcribe high amounts of RNA without inducing visible symptoms in the infected population (dos Santos et al., 2019) but, for example, infection with an iflavirus increases the susceptibility of *Spodoptera exigua* larvae to a baculovirus insecticide (Carballo et al., 2017). Interactions such as this among different species in a complex microbiome may explain the lower load of EVV-1 in *E. variegatus* individuals infected with Flavescence dorée phytoplasma (Abbà et al., 2017). The urgent need for more precise and sustainable pest management approaches to control phytoplasma diseases encourages the exploration of viruses as biocontrol agents able to compete with phytoplasma transmission and/or acquisition. The hypothesis that pathogen transmission may be suppressed or reduced by the concurrent vector infection with other microorganisms is supported by cases of West Nile virus vectored by mosquitoes co-infected with Nhumirim virus (Nouri et al., 2018) or with Wolbachia (Glaser and Meola, 2010), and by the cases of Dengue, Chikungunya, and Plasmodium transmitted by *Aedes* co-infected with Wolbachia (Moreira et al., 2009).

The biological information we provided here will be useful in the future to exploit EVV-1 for VIGS to regulate the expression of insect metabolic genes and, possibly, of genes involved in phytoplasma transmission. Also, an iflavirus-based vector will be a valuable tool to explore the role of potential phytoplasma effectors in the insect host. To reach this goal, preliminary results in the synthesis of an EVV-1 infectious clone have been obtained (Simona Abbà, personal communication) and RNAi mediated by injection of dsRNA molecules has been proven to work efficiently in *E. variegatus* with a long-lasting effect (Abbà et al., 2019). Elucidating the mechanisms of insect tolerance to iflavirus infection paves the way to conceptualize new anti-vectorial strategies to selectively control plant pathogen-transmitter Hemipteran insects.

## Author contributions

Concept & financing (LG, SA, CM, DB); Experimental design (LG, MR, CM); Execution (SO, AP, MR, MV, GM); Analysis (SO, MR, LG, CM); Publication (LG, MR, SA, MV, SP, DB, RB, CM). All authors read and approved the final manuscript. The authors declare no competing financial interests.

## Acknowledgments

The authors thank Elena Zocca for providing plants for insect rearing, Flavio Veratti for maintenance of insect colonies in Turin and Dr. Xavier Foissac for providing adults of *E. variegatus* from his laboratory colony, allowing us to set up a virus-free population.

This project has received funding from the European Union’s Horizon 2020 research and innovation programme under grant agreement No 773567. This research was also supported by Fondazione Cassa di Risparmio di Torino, Projects Siglofit (RF = 2016-0577) and FOotSTEP (RF = 2018-0678).

## Supplementary methods

### 2.5 RNA extraction, cDNA synthesis and EVV-1 qPCR assays

Total RNAs were extracted from samples of single *E. variegatus* adults, and pooled samples of i) 30 laid eggs, ii) three nymphs, iii) three dissected organs, iv) hemolymph collected from 10 insects, v) honeydew collected from 60 insects and vi) 500 mg of oat leaves. The samples were frozen with liquid nitrogen, crushed with a micropestle in sterile Eppendorf tubes, and homogenized in Tri-Reagent (1 mL for plant samples and 0.5 mL for all other samples). Samples were centrifuged 1 min at 12.000 g at 4°C and RNAs were extracted from supernatants with Direct-zol RNA Mini Prep kit (Zymo Research), following manufacturer’s protocol and including the optional DNAse treatment step. Concentration, purity, and quality of extracted RNA samples were analysed through a Nanodrop spectrophotometer (Thermo Scientific). For each sample, cDNA was synthesized from total RNA (1 μg) with random hexamers using a High Capacity cDNA reverse transcription kit (Applied Biosystems). Yields of reverse transcription reactions were estimated by reading a 1:10 cDNA dilution in a Nanodrop spectrophotometer (Thermo Scientific).

A multiplex qPCR assay was developed to detect and quantify virus presence in *E. variegatus* adults. To detect EVV-1, primers and FAM-labelled TaqMan probe were designed on the virus capsid 1 coding sequence (Evv1Cap1Fw 5’-GACCATTATCGCGCTAATG-3’, Evv1Cap1Rv 5’-AGTGCTCATCATAGGACA-3’, Evv1Cap1Probe 5’-FAM-ATTCTCGTAGCCAACTGCCAAAC-3’). To check that all the cDNA samples were properly amplifiable, the *E. variegatus* glyceraldehyde-3-phosphate dehydrogenase (GAPDH) transcript, was chosen as endogenous control. To amplify the insect GAPDH, the primer GapFw632 (5’-ATCCGTCGTCGACCTTACTG-3’ (Galetto et al., 2013)), was used with the newly designed primer GapRv819 (5’-GTAGCCCAGGATGCCCTTC-3’) and the internal HEX-labelled TaqMan probe GapEvProbe (5’-HEX-ATATCAAGGCCAAGGTCAAGGAGGC-3’). One μL of cDNA was used as template in a reaction mix of 10 μL total volume, containing 1X iTaq Universal Probes Supermix (Bio-Rad), 200 nM of each of the four primers and 300 nM of each of the two TaqMan probes. Each sample was run in triplicate in a CFX Connect Real-Time PCR Detection System (Bio-Rad). Cycling conditions were 95°C for 3 min and 40 consecutive cycles at 95°C for 10 s as denaturing step followed by 30 s at 60°C as annealing/extension step. In each qPCR plate, four serial 100-fold dilutions of pGem-T Easy (Promega) plasmids, harbouring the target virus and insect genes, were included to calculate the viral load. For both plasmid standard curves, dilutions included in plates ranged from 10^8^ to 10^2^ target copy numbers per μL and were prepared taking into account that 1 fg of plasmids harbouring EVV-1 and GAPDH gene portions contains 138 and 244 number of molecules, respectively. Dilution series of both plasmids were used to calculate qPCR parameters (reaction efficiency and R^2^). Mean virus copy numbers in amplified samples were automatically calculated by CFX MaestroTM Software (Bio-Rad) and used to express virus amount as EVV-1 copy numbers/insect GAPDH transcript.

## Supplementary Tables

**Supplementary Table S1.**
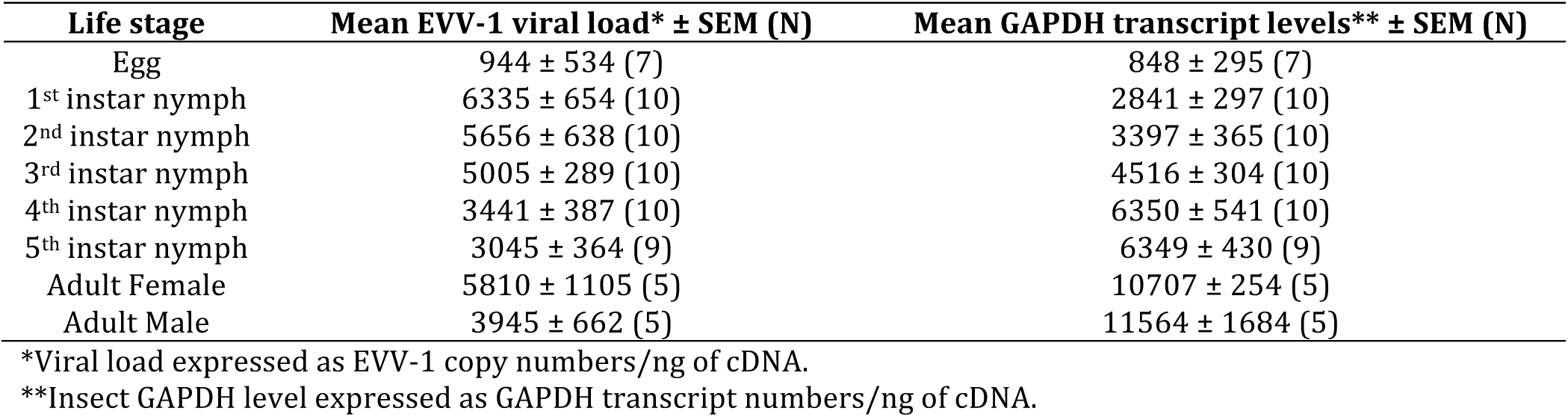
Presence and loads of Euscelidius variegatus virus 1 (EVV-1) in different insect life stages of the virus infected EvaTo rearing and corresponding mean transcript levels of insect glyceraldehyde-3-phosphate dehydrogenase (GAPDH). All samples were amplifiable.

**Supplementary Table S2.** Presence and loads of Euscelidius variegatus virus 1 (EVV-1) in different insect organs of the virus infected EvaTo population and corresponding mean transcript levels of insect glyceraldehyde-3-phosphate dehydrogenase (GAPDH). All samples were amplifiable.

| Organs | Mean EVV-1 viral load* ± SEM (N) | Mean GAPDH transcript levels** ± SEM (N) |
| --- | --- | --- |
| Malpighian tubules | 537 ± 185 (3) | 2665 ± 870 (3) |
| Salivary glands | 164 ± 36 (3) | 234 ± 41 (3) |
| Guts | 437 ± 66 (3) | 5118 ± 249 (3) |
| Ovaries | 3768 ± 1332 (5) | 5491 ± 1617 (5) |
| Testes | 16 ± 13 (3) | 132 ± 87 (3) |
| Hemolymph | 741 ± 290 (4) | 57 ± 21 (4) |

\*Viral load expressed as EVV-1 copy numbers/ng of cDNA.
\*\*Insect GAPDH level expressed as GAPDH transcript numbers/ng of cDNA.

**Supplementary Table S3.** Presence and loads of Euscelidius variegatus virus 1 (EVV-1) following injection into the EVV-1 free EvaBx population. Mean quantification cycles (Cq) obtained from insect glyceraldehyde-3-phosphate dehydrogenase (GAPDH) qPCR assay confirmed that all samples were amplifiable.

| Sample description | Sampling date (dpi, days post injection) | N° EVV-1 positive/ analysed samples | Mean EVV-1 viral load* ± SEM (N) | Mean Cq ± SEM (N) of GAPDH assay |
| --- | --- | --- | --- | --- |
| F0 (EvaBx adults injected with EVV1 extract) | 4 | 8/9 | 0.41 ± 0.15 (8) | 19.97 ± 0.64 (9) |
| | 10 | 10/10 | 1.37 ± 0.63 (10) | 20.20 ± 1.49 (10) |
| F1 (Progeny of injected EvaBx adults) | 60 | 16/16 | 2.40 ± 0.49 (16) | 17.41 ± 0.21 (16) |
| F2 | 120 | 10/10 | 2.53 ± 0.60 (10) | 16.91 ± 0.17 (10) |

\*Viral load expressed as EVV-1 copy numbers/insect GAPDH transcript.

